# Discovery of a sulfotyrosine-motif in the human TrkB extracellular domain required for agonist activation

**DOI:** 10.64898/2026.05.19.725324

**Authors:** David C. Briggs, Ryan T. Duffy, Sarah Ateaque, Sarah Maslen, Hema Nagaraj, Yves-Alain Barde, Peter S. DiStefano, Ronald M. Lindsay, Paul C. Armstrong, Chloe J. Peach, Neil Q. McDonald

## Abstract

The brain-derived neurotrophic factor (BDNF)-tropomyosin receptor kinase B (TrkB) signalling axis is a key effector of synaptic plasticity and neuroprotection. While TrkB activation is a major objective towards preventing dysfunction of the nervous system, it cannot be reached with exogenous BDNF administration given the unfavourable physiochemical properties of BDNF. In addition, BDNF also activates a tumour necrosis factor pathway by binding to the neurotrophin receptor p75. The TrkB agonist ZEB85 provides an alternative route to the selective activation of TrkB. We report here the structural basis for the interaction between human TrkB, and both ZEB85 and BDNF, and reveal that a sulfated tyrosine modification is indispensable for ZEB85 activation of TrkB signalling. Using structure-guided BDNF- and ZEB85-binding deficient TrkB mutants, we assessed their ability to sequester ligands from full-length TrkB in cultured human neurons. We found that the BDNF binding site extends into the extracellular juxtamembrane domain of TrkB but does not require the sulfotyrosine at residue 400 to activate TrkB. Together with biophysical analysis and AlphaFold modelling these results also explain how BDNF can displace ZEB85 from TrkB through an overlapping epitope. Our findings reveal unique features of TrkB, not present in the related neurotrophin receptors TrkA and TrkC, and suggest new directions to explore the role of sulfotyrosine in TrkB signalling and identify new TrkB-specific protein ligands.

**One Sentence Summary:** Investigation of the mechanism of action of TrkB agonist ZEB85 extends molecular understanding of TrkB activation.

## Introduction

Biological agonists against cell surface receptors have seen a renaissance in recent times, providing safe, developable and readily manufactured therapeutics (*1*). With biologics dominating the top selling drugs for 2025, this modality is set to continue expansion in the coming decade (*2*). Now that clinically approved solutions for delivery in the CNS have been found, a major growth area in neurology focused therapeutics is developing antibody biologics against CNS targets (*3*). As well as their potential transformational impact on neurological disease, antibody agonists also provide clean tools for discovery science by providing a route to elucidate their mechanism of action, imaging active receptor state location, uncovering conformational states, partial agonism and even unexpected modifications (*1*).

A key target for neuroprotection in neurodegenerative contexts is the tropomyosin-related <u>k</u>inase <u>B</u> (TrkB), a key signalling receptor for <u>b</u>rain-<u>d</u>erived <u>n</u>eurotrophic factor (BDNF)(*4*). TrkB is expressed widely in the CNS, and BDNF signalling through TrkB plays a crucial role in the synaptic regulation of brain physiology and pathology including neuronal differentiation (*5*), synaptogenesis (*6*), neuronal architecture (*7*), synaptic plasticity (*8*), depression (*9*) and neurodegeneration (*4*). Enthusiasm for BDNF therapy, despite showing promise in cellular and animal-models of neurodegeneration (*10*), has been tempered by inconsistent findings and complex underlying mechanisms of its action, and the poor bioavailability of BDNF. In addition to this, proBDNF and mature BDNF also show affinity towards the p75NTR receptor which has a quite distinct signalling pathway driving neuronal apoptosis (*11*). In contrast, recent attention has focused on agonists of TrkB in different formats such as small molecules (*12*, *13*) peptides (*14*), and antibodies (*15–18*). Among several reported TrkB antibody-based agonists is the full TrkB agonist ZEB85 (a fully human sc-Fv-Fc fusion protein) identified by coupling a function-based cellular assay to screening a combinatorial scFv library (*18*). Characterisation of ZEB85 using a human TrkB reporter cell line and BDNF-responsive GABAergic neurons, showed that ZEB85 is a high-fidelity agonist of TrkB with comparable signalling and transcriptional responses to BDNF regarding TrkB autophosphorylation (*18*). Furthermore, ZEB85 binding to TrkB was shown to be displaced by BDNF in a flow-cytometry based competition assay, suggesting at least some overlap in their respective TrkB binding epitopes (*18*).

The TrkB extracellular module shares the same domain composition as TrkA and TrkC, comprising a leucine-rich repeat domain (LRR), two consecutive Ig domains (IG1-IG2), and a more divergent extracellular juxtamembrane region (eJM) prior to the transmembrane helix. Insights from structures of the TrkA-NGF complex (*19*) and TrkB-NT4/5 heterodimer (*20*) have demonstrated the 2^nd^ IG domain of Trk receptors contains the key neurotrophin-binding determinants, with recent studies also suggesting the extracellular module is coupled to kinase activation through less well-defined regions such as the eJM (*21*).

The TrkB receptor extracellular module is known to be extensively glycosylated (*22*), but the heterogeneous nature of glycosylation can make other post-translational modifications hard to detect. Sulfotyrosine (sY or sTyr) is a relatively uncommon modification of extracellular proteins, first isolated in fibrinogen (*23*). Detection of sulfotyrosine residues has historically been quite difficult, as the sulfo group is not only smaller than the degree of heterogeneity typically observed in glycan modifications, but is also labile in strong acids that are used in Edman degradation and routine mass spectrometry protocols (*24*). Recent advances have led to protocols being developed to overcome this issue (*25*). Since no “eraser” activities have been found in mammals, sulfotyrosine is stable in most cellular environments (except for strong acids) and is considered an irreversible modification (*26*). In mammals, this tyrosine modification is “written” by one of two tyrosyl protein sulfotransferases (TPST1/2). TPST enzymes are single-pass type I transmembrane proteins with their catalytic domain located in the lumen of the trans-Golgi, able to transfer a sulfate group from a PAPS (3′-phosphoadenosine 5′-phosphosulfate) co-factor to the hydroxyl group of a tyrosine residue (*27*). Structure-function analysis of human TSPT1 and 2, and analysis of known sulfotyrosine-containing proteins shows a preference for acidic amino acids in close proximity to the target tyrosine residue(s) (*28*, *29*). Extracellular proteins within the complement and coagulation pathways are frequently modified by sulfotyrosine and are frequently found in or adjacent to protein-protein interaction sites where they contribute to the interaction surface and enhance binding affinity (*30*).

To understand the mechanism of action for ZEB85 (and other TrkB agonists) and its close mimicry of BDNF in terms of signalling outputs and transcriptional responses, we mapped the ZEB85 epitope on TrkB. We uncovered an unexpected sulfotyrosine modification in the TrkB epitope recognised by ZEB85, within an extracellular juxtamembrane region that is highly divergent between different Trk receptor isoforms. We determined ZEB85 and BDNF structures bound to their respective TrkB binding epitopes (including the first report of the structure of a TrkB^ECD^:BDNF complex) and used complementary biophysical and competition assays to reveal that both ZEB85 and BDNF interact with the TrkB extracellular juxtamembrane region.

## Results

### Mapping the ZEB85 epitope and discovery of a sulfotyrosine modification of TrkB eJM

To understand the molecular basis as to how ZEB85 mimics BDNF activation of TrkB, we first sought to map the ZEB85 epitope using purified proteins containing either the full human TrkB extracellular module (residues 32-432, TrkB^432^) or lacking the eJM (residues 32-383, TrkB^383^) (**Fig1A**) We used surface plasmon resonance (SPR) to measure binding of ZEB85 and found that ZEB85 bound TrkB^432^ with nanomolar affinity (K_D_ = 0.24 nM ± 0.08 nM), but showed no detectable binding to TrkB^383^ implicating the TrkB eJM as a harbouring a key binding epitope (**Fig1B,C**). As the eJM is predicted to be unstructured, we used a Western blotting approach to refine the epitope more precisely. We generated a series of finer TrkB truncations and transiently transfected constructs into HEK293 cells for analysis of their conditioned media for ZEB85 binding (**Fig1D**). We observed a dramatic loss of ZEB85 binding between the TrkB^402^ and TrkB^393^ truncations. We then examined human/rat chimeras in the eJM, taking advantage of the observation that ZEB85 activates human TrkB but is much less potent towards rodent TrkB (*18*). We identified four regions of substantial sequence divergence within the eJM and swapped these regions from the rat sequence into human TrkB. This gave generated human/rat chimeras TrkB_W381R, TrkB_YET, TrkB_WTTPT and TrkB_ADQSN **(Fig1D**). Only TrkB_WTTPT, corresponding to residues 400-405, showed no binding to ZEB85. Thus, both TrkB truncations and human/rat chimera experiments converged independently on residues 393-405 of TrkB as the key binding determinant of ZEB85 (**Fig1F**). Surprisingly a synthetic peptide spanning this region (residues 383-432 of TrkB) bound only weakly to ZEB85 with micromolar affinity (K_D_ 2.34 µM ± 0.91 µM), 4 orders of magnitude less than observed for recombinant TrkB^432^ (**Table S2**).

**Fig. 1.**
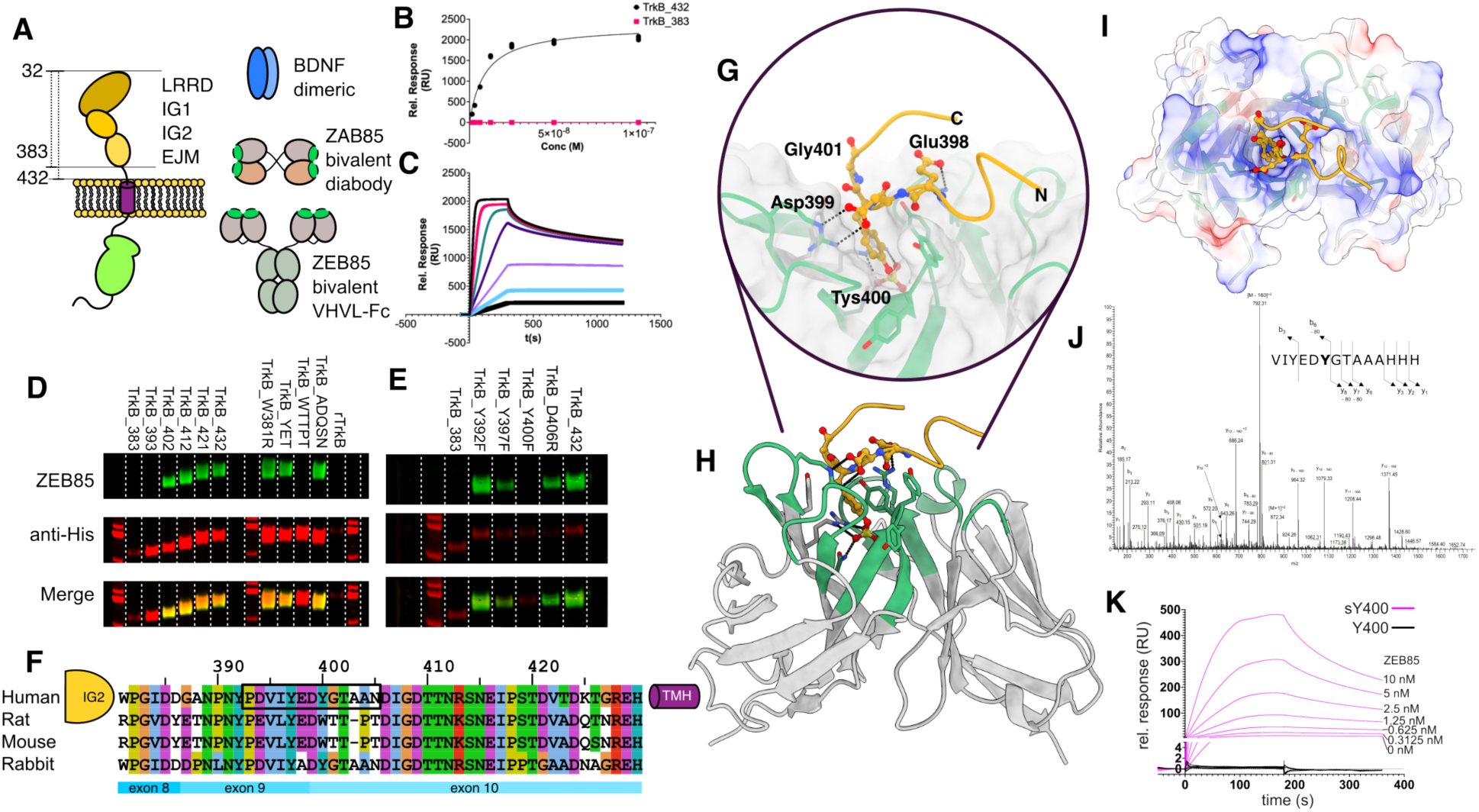
The full agonist ZEB85 recognises a sulfotyrosine-motif in the TrkB extracellular juxtamembrane domain. (**A**) Schematics of molecules described in this study (**B,C**) ZEB85 binds to TrkB^432^ but not TrkB^383^ with high affinity (K_D_ = 0.24 nM ± 0.08 nM (n=7)). (**D**) Representative western blots probing with ZEB85 (green) or anti-His tag control (red) showing that the ZEB85 epitope can be localized to residues 393-405 of TrkB, with (**E**) Tyr^400^ being a critical determinant. (F) comparison of selected TrkB orthologue eJM sequences between IG2 domain and the transmembrane helix (TMH). (**G,H**) Crystal structure of ZEB85 V_H_V_L_ bound to a motif in the TrkB eJM epitope showing close electrostatic complementarity and (**I**) a deep pocket for the sulfotyrosine moiety formed by the CDRs (**J**) Mass spectrometry evidence for sulfotyrosine at Tyr^400^ for recombinant human TrkB ectodomain digested with bromelain. (**K**) sTyr^400^-modified peptide reconstitutes nanomolar binding affinity to ZEB85 (KD = 0.52 nM ± 0.71 nM (n=3)), while unmodified peptide shows no detectable binding to TrkB.

We reasoned that the discrepancy between ZEB85 affinity for the synthetic peptide and recombinant TrkB^432^ may be due to a previously undetected post-translational modification in this region, and therefore explored that possibility by mass spectrometry. We discovered that our Expi293-expressed TrkB had a sub-stoichiometric +80 Da modification at residue Tyr^400^ (**Fig1J**), which could correspond to either a phospho- or sulfo-tyrosine modification. We prepared a panel of Y to F mutants within the ZEB85 epitope and showed that a Y400F mutation rendered the interaction between ZEB85 and TrkB undetectable, implicating Tyr^400^ modification as the critical determinant (**Fig1E**). Comparison of the amino acid sequence of TrkB surrounding the sulfotyrosine site (YED<u>Y</u>GTA) with other known sulfotyrosine-containing human proteins (**FigS1**) revealed that the acidic sequences flanking Tyr^400^ are similar to those found in known sTyr-containing proteins such as Nidogen-1 (DEDs<u>Y</u>DLA) and Complement C4 (s<u>Y</u>EDs<u>Y</u>Es<u>Y</u>D)(*30*) suggesting that this TrkB sequence could be a suitable substrate for tyrosine-O-sulfotransferases known to reside in the Golgi(*28*, *29*), whereas extracellular proteins phosphorylated by (for example) the secreted kinase VLK are modified at a G-pY-P motif (*31*). Based on these considerations, we considered the sulfotyrosine modification as more likely. We then used SPR to measure the affinity of ZEB85 for TrkB peptides (Biotin-linker-PDVIYED<u>Y</u>GTAAN) either with or without the sulfotyrosine modification. The modified peptide bound robustly to ZEB85 (0.52 nM ± 0.71 nM), recapitulating the tight binding measured for the recombinant TrkB^432^, whereas the unmodified peptide showed no detectable binding (**Fig1K**).

### ZEB85 recognises a linear sTyr-motif within the TrkB extracellular juxtamembrane region

To confirm the central role of sTyr^400^ and directly image the role of flanking residues in binding ZEB85, we determined crystal structures of a single chain Fv region of ZEB85 (“ZAB85”) either with or without the sTyr^400^-containing synthetic peptide (Acetyl-PDVIYEDsYGTAAN-NH_2_). Data processing and refinement statistics can be found in **Table S1**. The connecting linker between the V_L_ and V_H_ domains is disordered, but further refinement and biophysical analysis revealed that ZAB85 is a diabody with domain-swapped V_L_ and V_H_ domains(*32*) (see below and **FigS2**). The apo structure revealed a deep cleft comprised of aromatic and basic sidechains from CDRH1 & 3, with a non-CDR Arg^76^ and Asn^61^ sidechain from CDRH1 at the base of the pocket. In this apo structure, this pocket is partially occupied by a molecule of the buffer, HEPES with the sulfate-moiety pointing deep into the cleft.

The ZAB85-sTyr peptide complex structure showed unambiguous electron density for the core interacting region of the sTyr^400^-modified peptide. The structure reveals how the sTyr^400^ sidechain penetrates deep into the CDRH1/CDRH3 pocket (**Fig1G,H**), with extensive electrostatic interactions between the aforementioned non-CDR Arg^76^ and CDRH1 Asn^61^ residues at the base of the pocket and the sulfate moiety, with hydrophobic interactions between the aromatic group of sTyr^400^ and three tyrosine side chains present in CDRH3. The structure also reveals two salt bridges between Asp^399^ of TrkB and Arg^80^ (CDRH2), and of Glu^398^, and Arg^241^ (CDRL3)(**Fig1I**).

Each of the two copies of the TrkB/ZAB85 complex in the asymmetric unit exhibit a positive phi mainchain torsion angle at TrkB Gly^401^ indicating a key structural role for a glycine residue at this position. Amino-terminal to Glu^398^ in the TrkB peptide is a less well-ordered hydrophobic motif (V-I-Y). Tyr^397^ forms a hydrophobic interaction with Tyr^128^ (CDRH3), the sidechain of Ile^396^ approaches the Tyr^182^ sidechain, and the sidechain of Val^395^ approaches Tyr^181^. The two copies in the asymmetric unit diverge in this region, suggesting they are less important for the binding of ZEB85. Therefore, we conclude that the full TrkB epitope recognised by ZEB85 corresponds to Y-E-D-sY-G (TrkB^397-401^).

To validate our structural findings in cellulo, we assessed the effect of the Y400F substitution on human TrkB signalling activity in HEK293 cells. We measured TrkB activation by co-transfecting a genetically encoded FRET-based sensor of ERK activity using the extracellular signal-regulated kinase activity reporter, EKAR localised in the nucleus to track the real-time accumulation of nuclear phospho-ERK (pERK) downstream of TrkB activation (*33*). Both BDNF and ZEB85 activated wild-type TrkB even at low concentrations, with EC_50_ values of close to 0.2 nM for each ligand (pEC_50_ BDNF = 9.82 ± 0.26, pEC_50_ ZEB85 9.60 ± 0.17) (**Fig2A,B**). However, only BDNF was able to activate TrkB Y400F (**Fig2C**), while no ERK signalling was observed when TrkB Y400F was challenged with ZEB85 (**Fig2E,D**). This demonstrates the exquisite sensitivity of ZEB85 for sY-modified TrkB that is relevant for signalling in this cellular context. We concluded that the sulfation of Tyr^400^ is an absolute requirement for activation of human TrkB by ZEB85 but not for BDNF. Given our observation that ZAB85 formed a diabody, we tested whether the ZEB85 Fc domain fused to its scFv domain was dispensable, testing ZAB85 in the same real-time ERK assay with TrkB. ZAB85 exhibited a comparable EC_50_ to ZEB85 indicating the Fc domain was not crucial. (**Fig2F)**. However, when ZEB85 is reformatted as a full two-chain IgG antibody (ZIG85), its potency drops by several orders of magnitude (pEC_50_ ZIG85 7.37 ± 0.24 ;**Fig2F & FigS3**), suggesting that the relative spacing of the sY binding sites is key to efficient TrkB activation.

**Fig. 2.**
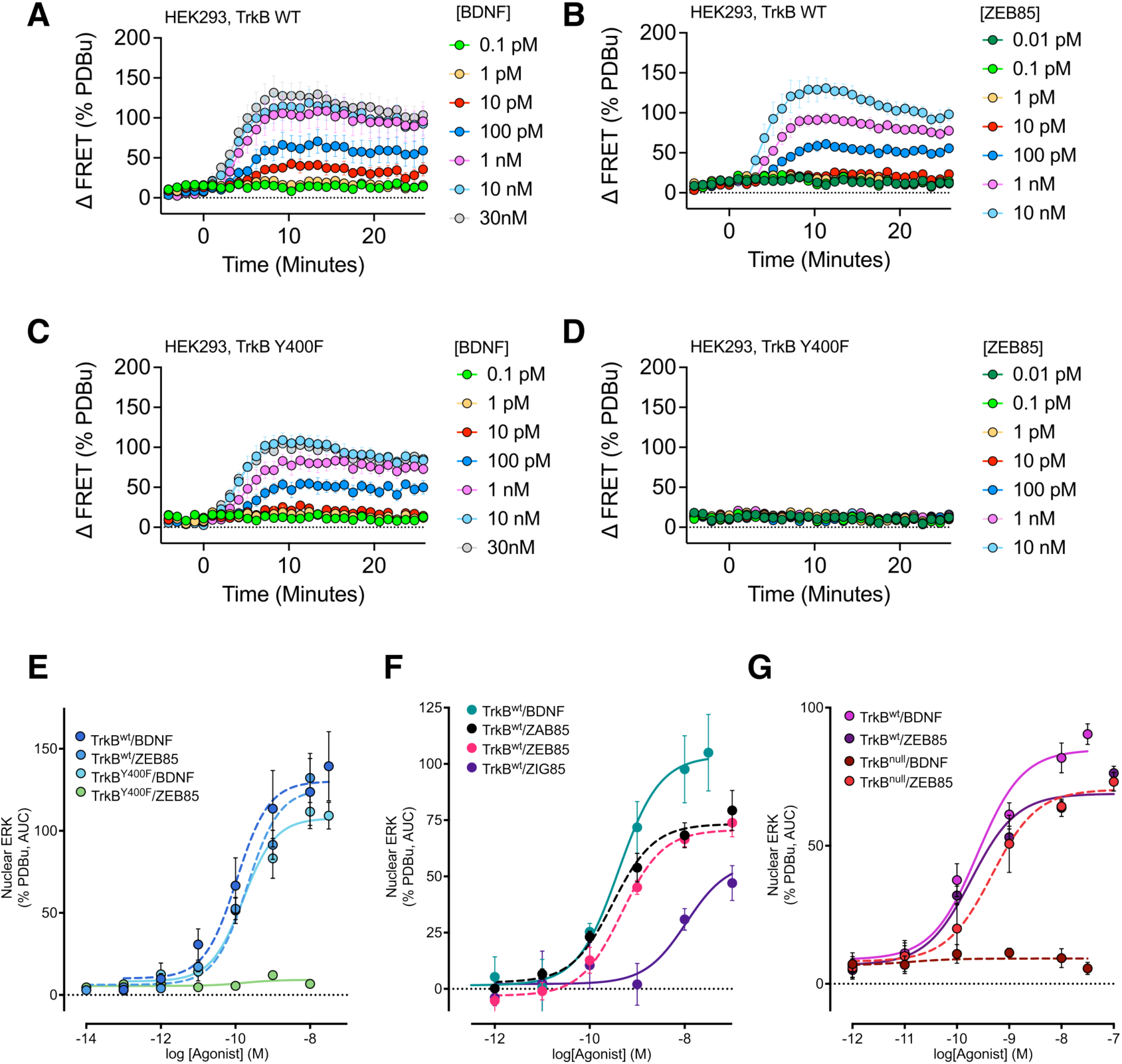
Cellular evidence that the Y400 sulfation site is crucial for ZEB85 agonist activation of TrkB signalling. (**A**) BDNF and (**B**) ZEB85 robustly activate wild-type TrkB signalling in transfected HEK293 cells using a nucEKAR FRET sensor measuring by nuclear accumulation of phospho-ERK. (**C**) BDNF fully activates TrkB Y400F signalling, but ZEB85 does not (**D**). (**E**) Comparison of EC50 curves from panels (A) – (D) for ZEB85 or BDNF stimulation of wild type or Y400F TrkB signalling. (**F**) Comparison of different ZEB85 formats as a Fab or full Ig antibody showing presentation of bivalent TrkB paratopes impacts agonism efficiency. Comparison of EC50 curves from examining activation of WT TrkB, or a TrkB^null^ (D298R/H299R/M379R) mutant that is deficient in BDNF binding, but still responds to ZEB85 **(G)**. Data from 4-5 independent replicates with triplicate wells. Mean ± SEM.

### ZAB85 is a diabody in solution and in the crystal lattice

To readily crystallize the ZEB85 V_H_V_L_ region containing the TrkB epitope, we removed the Fc region connected by a flexible linker in the ZEB85 (**Fig. 1A**). We anticipated that removal of the dimeric Fc region would produce a monovalent scFv (ZAB85). However, it became clear during this study that ZAB85 was as potent as ZEB85 in the FRET sensor assay for TrkB signalling. We investigated the monomeric/dimeric status of ZAB85 using mass photometry, which unambiguously revealed an in-solution molecular weight of double the expected monomeric molecular weight, consistent with a dimer even at low concentrations (**Fig. S2A**). Inspection of the packing in the crystal structure (indexed and scaled in point group 6/*mmm*) indicated that disordered peptide linker between the V_H_ V_L_ domains was unlikely to adopt a monomeric, side-by-side arrangement (either in the lattice or in solution), so we explored an alternative, lower point group and that might accommodate a domain swapped ‘diabody (*34*)’ (**Fig. S2B**). Ultimately, we settled on a point group of 1 (space group P1) and removed any imposition of symmetry. This reduced the overall resolution of the processed data, (as benefits of symmetry averaging were removed) but did allow us to extend the model for the disordered linker (**Fig. S2C**). This unambiguously demonstrated that ZAB85 was a diabody (*32*). Diabodies form when the V_H_ V_L_ domain connecting linker is too short forcing a non-covalent dimerization with another similar fragment, creating a stable molecule with two antigen-binding sites. We resolved two slightly different diabody structures in the asymmetric unit (**Fig. S2E,F** “Diabody AB” and “Diabody MN”), consistent with the two independent copies in the 6/*mmm* asymmetric unit. To further confirm the dimeric state of ZAB85 in-solution we performed size exclusion chromatography-coupled small angle X-ray scattering (SAXS) experiments and obtained data that are most consistent with the “AB” diabody (**Fig. S2D,G**) from the crystal lattice and provide further evidence for the diabody state of ZAB85. Based on the amino acid sequences of the ZEB85 and ZIG85 constructs, we conclude that the V_H_V_L_ region of ZEB85 also adopts the same diabody configuration (hence the comparable TrkB activation profile (**Fig. 2F)**, whereas in ZIG85, the extra domains and separate light and heavy chains mean this molecular adopts a classical IgG configuration with a much wider separation (134 Å ± 40 Å for IgG (*35*), compared with ∼50 Å for ZEB/ZAB85) of the CDR regions in the two arms of the antibody.

### Design of a BDNF-insensitive TrkB from the structure of a TrkB^383^-BDNF complex

The reported competition of ZEB85 and BDNF for TrkB binding (*18*) appeared at odds with the quite distinct molecular determinants predicted for BDNF within IG2 and shown here for ZEB85 in the eJM. Therefore, we sought to define a structure of BDNF bound to TrkB and to use this to design a “clean” TrkB mutant unable to respond to BDNF (that has yet to be reported). We prepared a complex using purified human TrkB^383^ and BDNF (residues 129-247, Icosagen) and determined its crystal structure (**Fig3A,B**). The diffraction data were anisotropic, but gave good quality density maps and a well-refined structure was obtained with good stereochemistry (**Supplementary Table 1**).

**Fig. 3.**
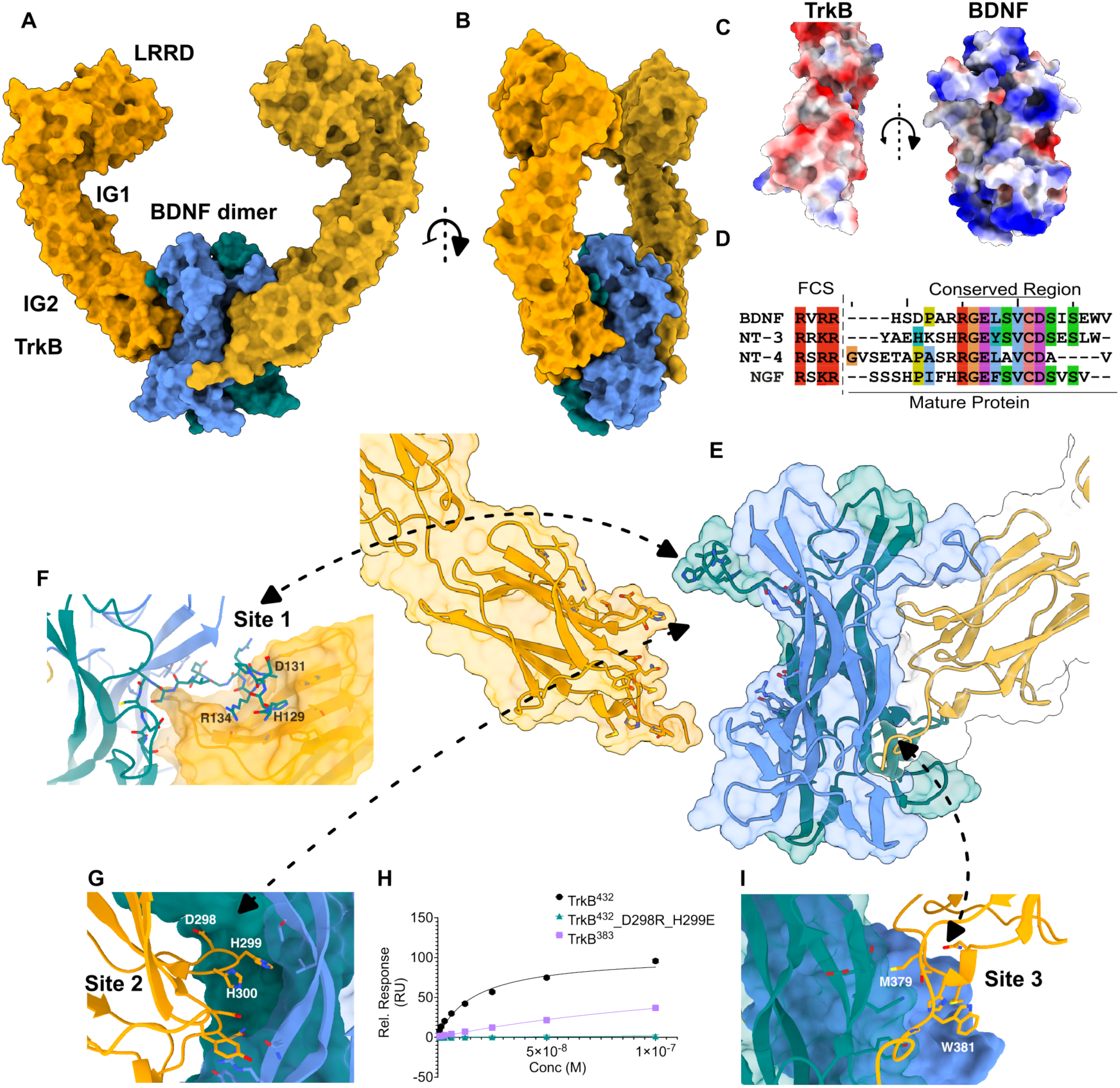
Overlap of BDNF and ZEB85 binding sites at the extracellular juxtamembrane region of TrkB. (**A,B**) Orthogonal views of the crystal structure of TrkB:BDNF complex with two main interaction sites. (**C**) Separated views for the TrkB:BDNF interaction, revealing electrostatic complementarity. (**D**) Comparison of mature neurotrophic factor amino terminal sequences. Separated views for TrkB and BDNF interaction revealing shape complementarity **(E)**. (**F**) The mature amino terminus of BDNF makes contacts with one face of TrkB IG2 (site 1). (G) Loops from IG2 of TrkB at the ‘waist’ of BDNF (site 2), making extensive contacts to both BDNF protomers. (**H**) SPR binding data show that TrkB^32-432^ binds BDNF more tightly than TrkB^32-383^, (TrkB^432^ KD = 9.8 nM ± 6.24 nM (n=3), TrkB^383^ KD = 35.7 nM ± 18.4 nM (n=3)) and that charge substitution mutants in one loop (D298R/H299E) abolish in vitro binding of BDNF to TrkB ectodomain. **(I)** Site 3 showing the packing of Met^397^ against a BDNF protomer.

The overall structure of the TrkB^383^- BDNF 2:2 complex is similar to that of TrkA^ECM^-NGF structure (*19*), with the dimeric BDNF ligand sandwiched between the 2^nd^ IG domains of the two TrkB ectodomains. There is a hinge motion between IG domains 1 and 2 of TrkA and TrkB that accounts for most of the gross structural differences between the two structures. BDNF forms a compact homodimer with a buried surface area of 1120 Å^2^ per monomer. The buried surface area between BDNF and one TrkB monomer is 1040 Å^2^, and the interfaces show extensive shape and electrostatic complementarity (**Fig3C,E**). The interface between TrkB and BDNF can be roughly segmented into three regions: an amino-terminal site (site 1) where the mature N-terminus from one molecule of BDNF packs against one face of TrkB IG2 (**Fig3F**), and a second site (site 2) where the residue 290s loop of IG2 packs against a shared interface composed of both BDNF protomers in the dimer (**Fig3G**), and a minor hydrophobic site at the membrane-proximal end of IG2 where Met^379^ interacts with a shallow hydrophobic pocket on one BDNF protomer (site 3, **Fig3I)**.

For site 1, the entire mature N-terminus of BDNF from His^129^ to Ile^138^ is ordered, revealing a key region that differs substantially from that of NGF (*36*), both in terms of sequence and structure when bound to their canonical Trk receptors (**Fig3D,F**). The first residue of BDNF after processing is His^129^ and both its sidechain and mainchain pack into a shallow groove on the face of TrkB IG2 formed by residues Phe^291^, Pro^304^, Thr^306^, His^343^, and the Cys^302^-Cys^345^ disulfide bond. We note that residues BDNF^129-135^ were disordered in heterodimeric structures of BDNF determined in the absence of TrkB receptor (*37*, *38*), suggesting that these amino acids become ordered on receptor binding. Comparison of our TrkB-BDNF structure with the TrkA-NGF structure suggests that site 1 confers specificity, with the amino-terminus of NGF/BDNF and 298-300 (TrkB numbering) loop of the receptor sterically hindering binding with mismatched ligands, supporting a hypothesis first put forward by Pattarawarpan and Burgess (*39*). We also note that the source of BDNF used in this study is the only commercial source with a native amino-terminal histidine, rather than an inauthentic initiator methionine or vector-derived leader sequence.

To validate these contacts and to design a TrkB variant insensitive to BDNF, we generated a panel of structure-guided mutations in TrkB probing residues at the interface with BDNF and assessed their affinities for BDNF by SPR (**FigS5**). While several mutants had a major impact on BDNF interaction, a double charge-reversal D298R/H299E mutant in the 290-loop of TrkB Ig2 domain exhibited the largest impact with no detectable interaction with BDNF in vitro (**Fig3H**). This mutant was then tested in the HEK293 nuclear ERK co-transfection assay, which unexpectedly retained activity. We therefore combined additional mutations to ultimately identify a triple mutant D298R/H299E/M379R (hereafter, TrkB^null^) that was completely unresponsive to BDNF in cellulo and gave no nuclear ERK signal but retained full activity for ZEB85 activation (**Fig2G, FigS4**). We concluded that full-length, cell surface TrkB exhibits a higher affinity for BDNF than the soluble TrkB ectodomain, perhaps through presentation of a preformed dimer mediated by interaction between the transmembrane helices as described by Kot *et al* (*40*), but that this triple mutation was sufficient to render TrkB insensitive to BDNF in cells. Thus, TrkB could now be altered selectively to respond to either BDNF or to ZEB85 through quite different separation-of-function mutations.

### BDNF binding surface with TrkB extends into the extracellular juxtamembrane

Our results revealed a clear distinction between ZEB85 and BDNF requirements for stimulation of TrkB with a sTyr^400^ modification. However, it was still possible that other elements in the eJM could be required for BDNF binding even if sTyr^400^ was not critical. We therefore investigated whether the initial series of TrkB truncations used to identify the ZEB85 epitope showed any impact on BDNF interaction by SPR. Results indicated a marked fall off in binding affinity for TrkB constructs ending between residue 393 and 402 (**Fig4A, Table S2**). TrkB truncations ending at or before residue 393 had apparent K_D_ values ∼40 nM, whilst truncated proteins ending after residue 402 showed apparent K_D_ values ∼10 nM. This suggested that an eJM binding contact for BDNF existed beyond the TrkB^383^ boundary used for structure determination. Additionally, we were unable to detect any significant difference between TrkB^432^WT (K_D_ = 9.8 nM ± 6.24 nM) and TrkB^432^Y400F (K_D_ = 15.3 nM ± 6.6 nM) binding to BDNF, consistent with the nuclear ERK FRET sensor experiments.

Attempts to crystallise a TrkB^402^-BDNF complex (with a longer eJM encompassing the sTyr modification) were not successful. We therefore used AlphaFold3 (*41*) to predict a potential binding site to include the eJM (**Fig4B,D, FigS6**). This gave a plausible model in good agreement with the SPR data, suggesting an extended interaction surface that continues as far as residue 405. Residues in this region have a predominantly acidic nature and therefore have good charge complementarity with the highly basic BDNF. Residues beyond this were not predicted to make contacts to BDNF, had low confidence pLDDT scores, and exhibited the characteristic “barbed-wire” appearance of a non-predictive AlphaFold result in this region (*42*).

To further validate the interaction of eJM with BDNF in a cellular context, we used a soluble TrkB ectodomain competition assay using mature human neurons derived from H9 embryonic stem cells that expressed endogenous full-length TrkB(*18*). The assay probed the ability of different soluble TrkB variants to sequester either BDNF or ZEB85 ligand and thereby block endogenous full-length TrkB activation as detected by tyrosine auto-phosphorylation (**Fig 4E,F**). For this assay, we first used two soluble variants of TrkB ECD to compete for BDNF or ZEB85 binding and prevent activation of cellular full-length TrkB. Whereas TrkB^432^ could sequester both BDNF and ZEB85 as predicted, preventing full-length TrkB activation (**Fig 4G**), TrkB^383^ was unable to sequester either ZEB85 or BDNF. For ZEB85, this can be explained by the loss of the sTyr^400^-motif, whereas the inability of TrkB^383^ to sequester BDNF agrees with the SPR data demonstrating that the eJM is required for BDNF interaction in this competition assay. To probe this further we used the same competition assay to examine the D298R/H299E charge reversal mutant that abolished BDNF binding to purified TrkB ectodomain in vitro and a human/rat chimera that removed the sY-motif of human TrkB (labelled WTTPT, **Fig 1D**) disrupting ZEB85 binding. Here there was a clear separation of function; BDNF was unable to bind the soluble TrkB D298R/H299E mutant resulting in no competition with full-length TrkB and thus activation. In contrast, ZEB85 was captured by this mutant blocking full-length TrkB activation. Conversely, TrkB WTTPT could compete with BDNF blocking full-length TrkB activation, whereas the same mutant could not bind ZEB85 and therefore was unable to block full-length TrkB activation (**Fig 4H**).

**Figure 4:**
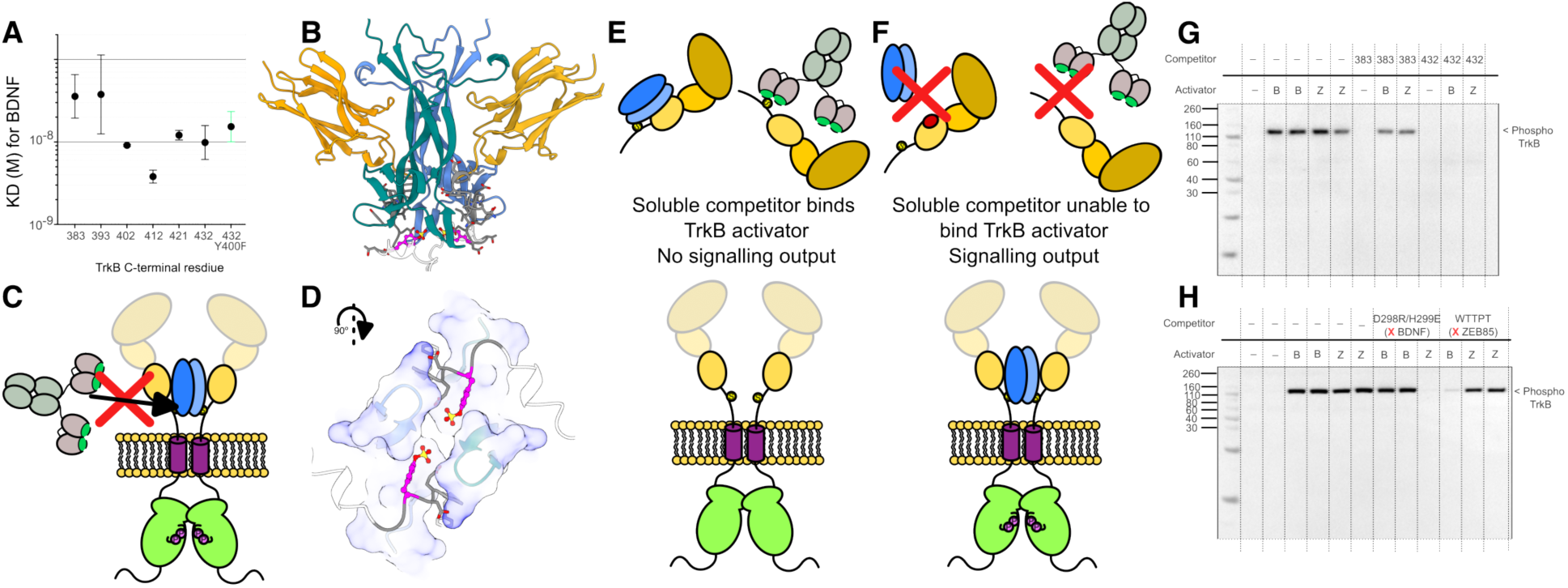
**(A)** Truncation series of the TrkB eJM reduces the affinity of TrkB for BDNF with a sharp K_D_ jump between TrkB^32-393^ and TrkB^32-402^ (n=3) **(B,D)** AlphaFold3 models align with our biophysical and cellular binding data, revealing an extended TrkB/BDNF interface with good electrostatic complementarity in which residues 383-402 of the eJM interact with and tuck under the membrane proximal surface of BDNF (for PAE plots & pLDDT colouring, see **FigS6**). **(C)** In this AF3 model, the ZEB85 epitope is sterically occluded by BDNF binding, explaining the observed competition. (**E,F**) Competition assay in H9 neurons between soluble recombinant TrkB ectodomain variants and endogenous full-length TrkB. **(G)** TrkB^32-383^ lacking the eJM cannot compete for either ZEB85 or TrkB activation leading to full-length TrkB activation. TrkB^32-432^ with the eJM motif sequesters both TrkB and BDNF preventing full-length TrkB activation. (**H**) A TrkB mutant selectively deficient in either BDNF (D298R/H299E) or ZEB85 (sulfotyrosine-motif deficient) binding cannot sequester the corresponding ligand and can’t block full-length TrkB activation.

As our AF3 model also includes the ZEB85 epitope in the extended binding site, this explains the observed displacement of ZEB85 from TrkB-expressing HEK293 cells by BDNF detected by FACS sorting(*18*). From these combined results, we conclude that the TrkB eJM contributes binding determinants for both BDNF and ZEB85 interaction, but only ZEB85 has an absolute requirement for the sulfotyrosine motif (**Fig 4C**).

## Discussion

TrkB agonist antibodies have long been recognized as promising tools to selectively activate TrkB (reviewed in (*43*)). Several potent antibodies have been described and are being pursued as potential therapeutics in neurodegeneration including 29D7, AB2 and AB20, mAb1D7, ZEB85, MM12, and AS86 (*16–18*, *45–47*). Interestingly, the epitope of MM12 has also been mapped to the TrkB eJM region, although the precise epitope remains unknown. However, the structural basis for their potent agonism is largely unknown despite how closely agonists such as ZEB85 can mimic TrkB-mediated signalling and transcriptional responses (*18*). Here, we identify a short linear sulfotyrosine-motif within human TrkB as the critical epitope critical for ZEB85 activation of TrkB. Using structure-guided separation of function mutations, we also show that ZEB85 and BDNF have overlapping TrkB binding sites within the extracellular juxtamembrane region.

### Overlap of BDNF and ZEB85 binding site at the TrkB extracellular juxtamembrane region

In spite of their overall sequence similarities, the 3 neurotrophin receptors TrkA, TrkB and TrkC show very different functional behaviours in vivo, including the induction of cell death from TrkA and TrkC, but not TrkB, in the absence of their respective ligands (*48*). The 4 mammalian neurotrophin ligands BDNF, NT4, NGF and NT3 distinguish the Trk receptors with varying degrees of selectivity. The extracellular juxtamembrane region of TrkB is a major source of sequence divergence among the Trk receptors, where the eJM segments of TrkA and TrkC are markedly shorter than TrkB and devoid of tyrosine residues. Our finding that BDNF and ZEB85 binding sites overlap at the eJM is supported by competition experiments (*18*) but had not been previously demonstrated (**Fig5**). A closer analysis of the eJM region by directly measuring BDNF affinity constants and using a competition assay for soluble TrkB ECM ligand sequestration, point to this region having a necessary but not sufficient ancillary role in BDNF binding. This then explains how ZEB85 with a nanomolar affinity could be competed off TrkB by BDNF interaction with a picomolar affinity by engaging the eJM segment. An important implication of this finding is that it extends the TrkB/BDNF binding site beyond the IG2 domain and towards the membrane. Such an extended interface had been originally proposed by Ibáñez and colleagues in a study detailing the generation of a pan-neurotrophin-1 (PNT-1) that could activate TrkA, TrkB and TrkC (*49*). Even a single chimeric molecule using an NT-3 backbone and BDNF variable region V (motif Ser^222^-Lys^223^-Lys^224^-Arg^225^-Ile^226^-Gly^227^) was able to enhance TrkB interaction and increased the survival of nodose neurons (*49*). In the crystal structure presented here, this basic region V motif in BDNF is fully accessible and predicted by AlphaFold to engage the acidic eJM of TrkB, consistent with our findings. One limitation of this study is the affinities are measured using soluble TrkB proteins rather than on the membrane. However, the competition assay does directly compare the ability of full-length TrkB to compete with soluble forms. Equally, using a FRET-based sensor to indirectly measure TrkB activation also confirm our in vitro findings. Both approaches could be used to identify directly the BDNF residues contacting TrkB eJM. Reported findings for other TrkB agonists have implicated binding determinants within the eJM but have not mapped the site to a specific motif such as the sulfotyrosine motif reported here (*46*). It is possible some of the reported TrkB agonist antibodies target sTyr^400^ and with the tools reported here these can now be readily tested. Other neurotrophins may also utilize their canonical Trk receptor eJM region. For example, a recent report for the NGF^painless^ mutant R100W proposes a TrkA eJM acidic contact site is impacted by this mutation, perturbing PKCγ signalling and uncoupling nociception signalling (*50*). The importance of the eJM in other RTKs has also been demonstrated, with different splice variants of ErbB4 and FGFR2 varying in their eJMs, which in turn leads to different signalling outputs (*51*) or ligand binding (*52*) respectively.

Additionally, our results that show that the ZIG85 variant, with different spacing of the paratopes, leads to a considerable loss in TrkB activation efficiency (**Fig. 2F**).This finding might have broader implications and suggests that refactoring antibody agonists into scFv-Fc scaffolds be brought into the toolkit of agonist antibody development alongside Fc-(*53*) and disulfide-engineering (*54*). Based on our structures, BDNF and ZEB85/ZAB85 have much smaller paratope-paratope distances of ∼30 Å (BDNF) and ∼50 Å (ZAB85), whereas mean paratope-paratope distances in IgG are ∼130 Å ± 40 Å (**Fig.5**) (*35*). Therefore we conclude that, much like other receptor tyrosine kinases (*55*), the spacing enforced by the TrkB domains closest to the membrane is of great import when considering efficient signalling output.

**Figure 5:**
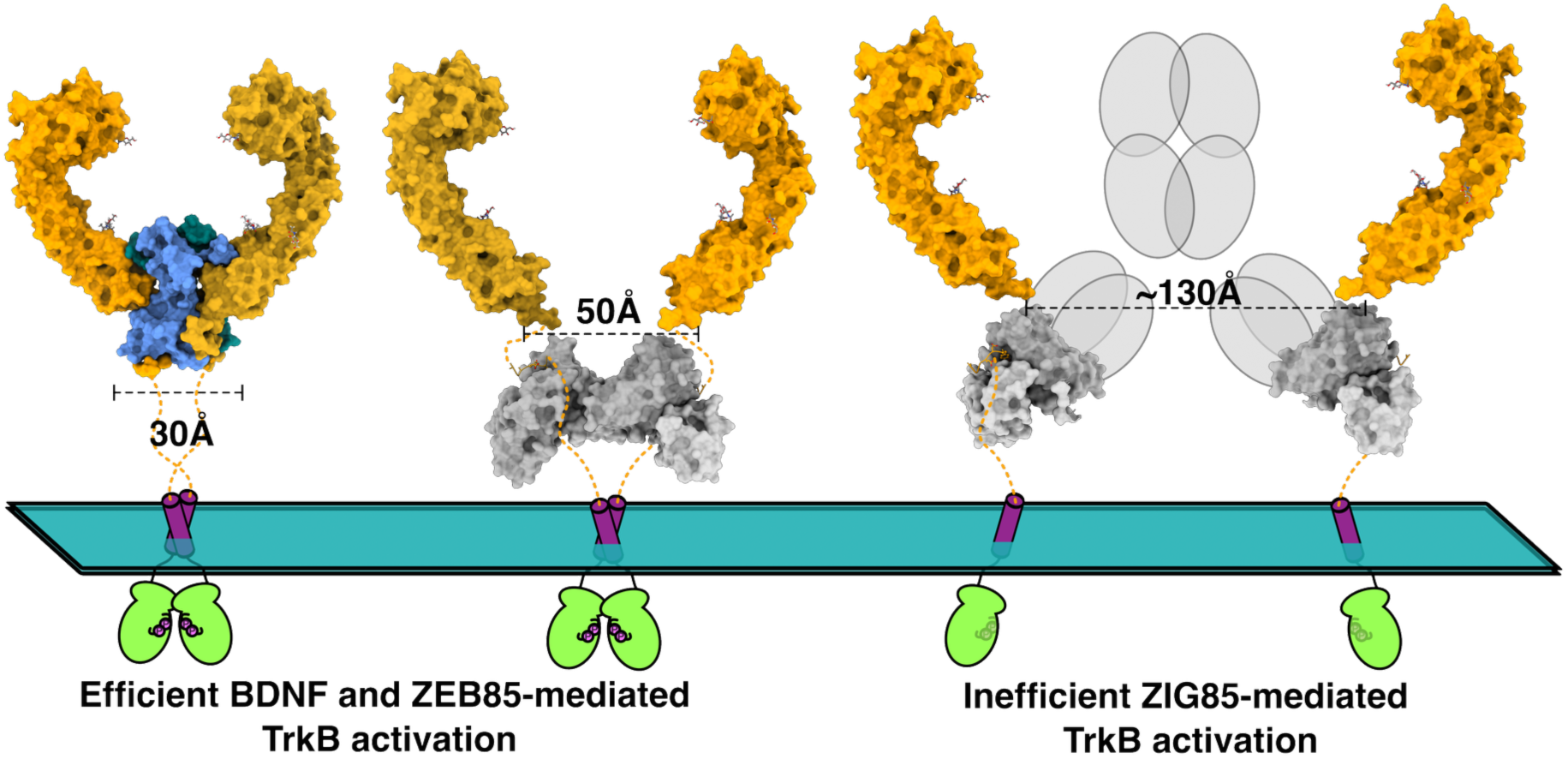
Schematic model of observed geometric constraints on TrkB activation. BDNF (blue) and ZEB85/ZAB85 (silver) permit close association of TrkB protomers (orange) (∼30-50 Å separation of TrkB protomer eJMs), leading to efficient activation. ZIG85 holds the TrkB monomers further apart (∼130 Å) leading to less efficient formation of a productive signalling complex.

### A potential physiological role of the sulfotyrosine motif unique to TrkB

To our knowledge, this is the first description of a receptor tyrosine kinase with a sulfotyrosine modification, which we have discovered by a combination of mass spectrometry, homology to known TPST substrates, and by virtue of its extracellular location. The functional role of this enigmatic modification is not yet clear, and may not be easy to explore in the most commonly used animal models given the absence of this tyrosine residues in the corresponding mouse and rat sequences. Given the affinity of ZEB85 for this modification, we propose the TrkB sulfotyrosine-motif could provide a protein interaction platform for physiological ligands, potentially analogous to intracellular SH2 domains for phospho-tyrosine (*56*). TrkB sulfotyrosine-partners are likely in neuronal contexts that could functionally impact TrkB signalling, stability or trafficking. Precedents exist for extracellular protein sulfotyrosine modification contributing to protein ligand-receptor interactions. For example, complement C5a binding to the C5a receptor is enhanced by three orders of magnitude by the presence of a pair of sulfotyrosine residues (*57*). Equally, multiple chemokine receptors are known to be sulfated at tyrosine sites within their amino-termini to influence the affinity of cognate chemokine interactions (*58*). The functional role of the motif is likely specific to mammalian TrkB since the motif is conserved in all major mammalian families apart from rodent TrkB sequences, which responds only weakly to ZEB85. Equally, the motif is unique to TrkB, as TrkA and TrkC lack any tyrosines in their respective eJM regions, preventing exposure to such a modification during Trk receptor maturation and passage through the trans-Golgi network. It is known that the sTyr modification is frequently sub-stoichiometric and it is possible that variation of the frequency of modification would allow for fine-tuning of sTyr-dependent TrkB functions, as is seen for CCR5(*59*). It is also known that a splice variant of Chicken TrkB lacks exon 9 (which encodes the region of the eJM containing the sTyr motif) has an increased specificity for BDNF over NT3, providing more evidence that alteration of this region does have a role in fine tuning TrkB:neutrophin engagement (*60*). For TrkB, there are a number of reported binding partners for the TrkB extracellular domain that might potentially exploit this motif, such as SLITRK5 (*61*) or the mGlu2 (*62*) or mGlu5 receptors (*63*, *64*). However, further work will be required to investigate if there are physiological roles and partners for the sTyr modification of TrkB.

Overall, our data point to a clear role that the TrkB eJM plays in recognition by BDNF and ZEB85, proposed to occur through a previously undetected and acid-labile sulfotyrosine modification. This study highlights how antibody agonists can reveal previously uncharacterized features and motifs relevant to endogenous ligand recognition. This can open up new hypotheses and point to new mechanisms and biology to explore, such as cross talk between TrkB and other synaptic receptors, or modulation of TrkB trafficking. Our study also highlights the increasing need in this post-genomic and post-AlphaFold era to detect, characterise and understand the contribution of underexplored post-translational modifications, such as sulfotyrosine, in human health and disease (*65*).

## Materials and Methods

### Protein production and purification

All proteins in this study were expressed in and secreted from Expi293 or Expi293 GntI- cells (Gibco) growing in Freestyle293 media (Gibco), transiently transfected using a PEI-based method as described in Pulido et al (*66*). A codon optimised human TrkB construct 1-383 (including native signal sequence 1-31) was synthesised by Geneart, with a series of mutants to remove non-conserved N-linked glycans to make the protein product more amenable to structural study by X-ray crystallography. This was subcloned into pcDNA3.1 vector with a carboxy-terminal hexahistidine-tag (6xHis-tag) to facilitate purification and detection. The construct for the full ECD of TrkB (1–432) was created by inserting the native sequence for TrkB residues 384-432 into this vector using Gibson assembly, retaining the 6xHis-tag. Mutations were introduced using Q5-polymerase site directed mutagenesis protocol (New England Biolabs). All TrkB extracellular proteins were harvested from the conditioned media with Ni-NTA Sepharose (Qiagen) washed with Tris-buffered saline (TBS) containing 10 mM imidazole and eluted using TBS with 500 mM Imidazole.

ZEB85 (V_H_V_L_-FC) was similarly generated by Geneart and subcloned into pcDNA3.1 and harvested using protein G Sepharose, and a low pH elution (0.2 M Glycine pH 3.5) step which was then immediately neutralised using 1 M Tris pH 9.0. A construct for the smaller ZAB85 (V_H_V_L_-3C-StrepII) was created using Gibson assembly, by inserting the coding sequence for the secretion sequence and V_H_V_L_ region into a pcDNA3.1 vector containing a C-terminal 3C cleavage site and a single StrepII-tag for purification. ZAB85 was harvested from conditioned media using streptactin Sepharose (IBA), washed with TBS, and then cleaved from the resin using GST-3C protease overnight at 4°C. The GST-3C was then removed from the eluate using GST-Sepharose (Cytiva).

All proteins were subjected to a final size exclusion chromatography polishing step using a Superdex 200 increase column (Cytiva). This material was then aliquoted and snap frozen in liquid nitrogen and stored at -70°C until needed. Mature BDNF was a generous gift from Professor Mart Ustav (Icosagen, Estonia).

### Surface plasmon resonance

All surface plasmon resonance experiments were performed on a Biacore S-200, in HEPES- or Tris-buffered saline with 0.005 %(v/v) P-20 surfactant. Chip chemistry, ligand, and analyte are detailed in table S2. Note that for BDNF - TrkB interactions, when the TrkB was the immobilised ligand and BDNF the analyte, the SPR sensorgrams were uninterpretable due to the approximately zero off rate consistent with the reported picomolar affinity of this interaction (*67*, *68*). We found that when the BDNF was immobilised, the apparent KD was reduced to ∼nM (presumably to stearic hinderance or non-optimal binding-site presentation on the sensor chip) giving an off-rate within the capabilities of the instrument and allowing us to compare different TrkB variants for BDNF binding.

### Western blotting to determine ZEB85 epitope

All gels were run using BOLT gels (Lifetec) in a MES running buffer. Transfer to nitrocellulose membranes was done using an iBlot2 or iBlot3 semi-dry transfer device. A blocking buffer of TBS + 0.1 % Tween (TBS-T) + 1 % BSA + 5 % milk powder was used for all primary antibodies. Incubation with primary antibodies was done overnight at 4°C. The membranes were then washed 3x10minutes with TBS-T before incubation with an appropriate LiCor secondary antibodies in fluorescence blocking buffer (Antibodies online) for 1 hour at room temperature. The membranes were then washed 3x10min with TBS-T again before visualising the using a LiCor Odessey scanner (LICORbio).

### Detection of sulfotyrosine by mass spectrometry

Purified TrkB^32-402^ was reduced and alkylated in solution and digested with bromelain at 1:20 ratio. The resulting peptides were analysed by nano-scale capillary LC-MS/MS using an Ultimate U3000 HPLC (ThermoScientific Dionex, San Jose, USA) to deliver a flow of approximately 300 nL/min. A C18 Acclaim PepMap100 5 µm, 100 µm x 20 mm nanoViper (ThermoScientific Dionex, San Jose, USA), trapped the peptides prior to separation on a C18 Acclaim PepMap100 3 µm, 75 µm x 250 mm nanoViper (ThermoScientific Dionex, San Jose, USA). Peptides were eluted with a 60 minute gradient of acetonitrile (2 % to 50 %). The analytical column outlet was directly interfaced via a nano-flow electrospray ionisation source, with a hybrid quadrupole orbitrap mass spectrometer (Q-Exactive Orbitrap, ThermoScientific, San Jose, USA). Data dependent analysis was carried out, using a resolution of 70,000 for the full MS spectrum, followed by ten MS/MS spectra. MS spectra were collected over a m/z range of 350–1800. MS/MS scans were collected using a threshold energy of 27 for higher energy collisional dissociation (HCD). LC-MS/MS data were searched against an in-house protein database using the Mascot search engine programme (Matrix Science, UK)(*69*). Database search parameters were set with a precursor tolerance of 10 ppm and a fragment ion mass tolerance of 0.15 Da. No enzyme was chosen for bromelain and variable modifications for oxidized methionine, carbamidomethyl cysteine, and sulphated serine, threonine and tyrosine. MS/MS data were validated using the Scaffold programme (Proteome Software Inc., USA)(*70*). All data were additionally interrogated manually.

### Synthesis and purification of sY containing peptides

The neopentyl-protected tyrosine-sulfated peptides were synthesised on a 0.1mmole scale, using solid-phase peptide synthesis on a ResPep Automated Peptide Synthesiser (CEM) using 5 equivalents of N(a)-Fmoc amino acids and 4.9 equivalents of HATU (Hexafluorophosphate Azabenzotriazole Tetramethyl Uronium) as the coupling reagent and 10 equivalents of Diisopropylethylamine (DIPEA). The Fmoc deprotections were carried out on the synthesiser using 20% piperidine in DMF (Dimethylformamide) containing 1% formic acid. Standard Fmoc protected amino acids were used, along with Fmoc-Tyr(S03nP)-OH (CAS 878408-63-0). All amino acid were double coupled.

Following chain assembly, peptides were cleaved from the resin and protecting groups removed by addition of a 10ml cleavage cocktail (95% TFA (trifluoracetic acid), 2.5% H2O, 2.5% TIS (triisopropylsilane) and 2.5% EDT (ethanedithiol) if the peptide contains cys or met residue. After 4h, the resin was removed by filtration, and the peptides were co-precipitated with 40mls of cold diethyl ether on ice. The peptides were isolated by centrifugation, then dissolved in 10mls H2O and freeze dried overnight. Following lyophilisation, the neopentyl (nP) group was removed by dissolving the peptide in a minimal amount of DMSO (40mg in 300ul). The solution was diluted to 1.5ml with 2M NH4OAc and incubated at 37 °C for 6 h. The reaction was monitored by using LC-MS.

After deprotection, the peptides were purified in 40mg batches on a C8 reverse phase HPLC column (Agilent PrepHT Zorbax 300SB-C8, 21.2x250 mm, 7 m) using a linear solvent gradient of 0-20% B over 40mins at a flow rate of 8 mL/min. A binary solvent system [A: H2O /0.08% TFA/1% MeCN (acetonitrile) and B: MeCN /0.08% TFA] was used. The peak fraction was analysed by LC–MS on an Agilent 1290 LC-MSD (Agilent Poroshell 120 EC-C18 2.7um, 3.0x30mm column), using a linear gradient of 5-60% B over 8.5 min at a flow rate of 0.425 mL/min.

### Crystal structure determination of ZAB85 alone and bound to TrkB^393-405^ eJM peptide

For ligand-free structure determination, ZAB85 was gel-filtered in HEPES-buffered saline and then diluted to 1:3 with milliQ water and concentrated to 5 mg/ml. Crystals grew in 17 % PEG3350, 0.24 M CaCl_2_, 0.1 M BisTrisPropane, pH 6.8, and were then cryocooled for data collection using 20 % ethylene glycol as a cryoprotectant. Data were collected to 1.88 Å resolution at beamline I04, Diamond Light Source. Diffraction images were indexed, integrated and scaled using DIALS (*71*) software. Phases were estimated using Phaser (*72*) and a molecular replacement search model of the VHL portion of 5YAX (*73*), with the CDR residues removed to reduce the influence of phase bias in the resulting maps. CDRs were readily traced and built into the resulting maps using Coot (*74*).

For structure determination of ZAB85 bound to a synthetic TrkB^393-405^ peptide containing the sTyr^400^ modification, the ZAB85 was gel filtered in Tris-buffered saline instead of HEPES buffered saline. Again, protein was diluted 1:3 with milliQ water and concentrated to 5 mg/ml. Ten-fold molar excess of solid peptide was dissolved in the protein solution prior to crystal screening. Crystals were grown in 20 % PEG8k, 0.2 M MgCl_2_, 0.1 M Tris pH 8.5, and were then cryocooled for data collection using 20 % ethylene glycol as a cryoprotectant. Data were collected to 2 Å resolution at beamline I24, Diamond. Diffraction data processing was performed as described for ZAB85 alone. Phases were estimated using Phaser (*72*) and a molecular replacement search model of the ZAB85 apo form, with the CDR residues removed to reduce the influence of phase bias in the resulting maps. The CDR residues and bound peptides were readily traced and built into the resulting maps. Final data processing and model statistics are given in Supplementary table 1.

### Crystal structure determination of TrkB^32-383^ bound to mature human BDNF

TrkB 1-383 (32-383 mature protein) was purified from Expi293 GntI- cells to produce homogenous N-linked glycan modifications. This protein was harvested from conditioned media using Ni-NTA Sepharose, and then further purified using size exclusion chromatography in buffer consisting of 20 mM Tris-HCl, pH 8.0, 100 mM Potassium Acetate. The material was then concentrated to 8 mg/ml and mixed with an appropriate amount of human BDNF at 1 mg/ml to yield a 1:1 molar ratio. This material was then used directly for crystal screening. Crystals grew from the Morpheus screen (*75*) in 30 % PEG4k/glycerol mix, 0.1 M Bicine/Tris buffer at pH 8.5 and 0.12 M Morpheus ethylene glycol mix. These crystals diffracted anisotropically to 3Å - 4.5Å at I04, Diamond Light Source. Data was processed using DIALS (*71*) – the unit cell parameters were initially identified as C222(1), 57.54 Å x 179.93 Å x 183.81 Å. The data were phased using the TrkA/NGF structure, PDB 2ifg (*19*), divided into 4 individual search models (TrkA LRRD, TrkA IG1, TrkA IG2, NGF) using phaser (*72*). Refinement stuck at an RFree of around 40 %, and so we hypothesised that the space group assignment may be incorrect. Zanuda (*76*) indicated that the true space group was C2. After reprocessing the data in C2 (with β = 90.04°), re-refinement gave a drop in RFree to ∼32 %. The data were further processed using Staraniso (*77*) (global phasing) to help ameliorate the effects of the anisotropy. Final data processing and model statistics are given in Supplementary table 1. All structural figures were generated using ChimeraX (*78*)

### TrkB signalling FRET sensor assay for nuclear-ERK activity

We used a genetically encoded, FRET-based sensor of ERK activity (the extracellular signal-regulated kinase activity reporter, EKAR) to assess TrkB signalling in transfected HEK293 cells(*33*). HEK293 cells plated (25,000 cells per well) in black 96-well plates pre-coated with poly-D-lysine were serum-starved following 24h and transfected with nuclear EKAR CFP YFP (40 ng/well; Addgene, #18681) and either FLAG-TrkB (WT) or FLAG-TrkB (Y400F mutation), SnapTag-TrkB (WT) or SnapTag-TrkB^null^ (D298R/H299E/M379R) mutation (20 ng/well) using polyethyleneimine (PEI, 6:1) diluted in PBS solution. Following 48 h, cells were incubated in Hank’s Buffered Saline Solution (HBSS)/0.1% Bovine Serum Albumin (BSA) and stimulated with increasing agonist concentrations either BDNF (0.1 pM-30 nM) or ZEB85 (0.01 pM-1000 nM) or ZAB85 (1 pM-1000 nM) or ZIG85 (1 pM-1000 nM). Phorbol 12,13-dibutyrate (PDBu, 10 µM) was used as a positive control and the buffer solution HBSS/0.1 % BSA was used as a vehicle treatment. Fluorescent emissions (CFP = 480 nm and YFP = 530 nm) were recorded (PHERAstar) over a period 26 minutes, with ligand addition occurring at t = 5 mins. FRET ratio was calculated (530nm/480nm), which provided the “raw data” for analysis. Data was then baseline corrected to the first 5 reads (pre-ligand addition) and then baseline corrected a second time to the vehicle. The data was then normalised to the maximal response of the positive control (PDBu, 100 %) or vehicle (0 %) and expressed as a percentage. Concentration-response data was fit using non-linear regression analysis as log vs. response (three parameters) to determine EC50 values. Data are presented as mean ± SEM and analysed using GraphPad Prism (10.1.2).

### Competition assay using iPSC-derived H9 neurons expressing endogenous full-length TrkB

Neurons differentiated from human ES cells (H9 clone) were prepared as described in(*79*). These neurons were incubated with the appropriate ligand for various lengths of times as indicated. Prior to lysis, the cultures were washed with PBS, and subsequently lysed with 70 μl of RIPA lysis buffer: 50 mM Tris- HCl, pH 7.4, 150 mM NaCl, 1 mM EDTA, 1% Triton-X-100, 0.2% sodium deoxycholate, 0.1% SDS and supplemented with a cocktail of protease and phosphatase inhibitors (Sigma-Aldrich) at 1:100 dilution containing 100 mM 1,10-Phenanthroline, 100 mM 6-aminohexanoic acid, 10 mg/ml aprotinin and 2 mM sodium orthovanadate. Cell lysates were kept on ice for 10 min, then centrifuged for 15 min at 15,000 rpm and the supernatant transferred to a new tube prior to western blot analysis, or storage at -80°C.4 X NuPAGE loading buffer (containing 0.666 g Tris HCl, 0.682 g Tris Base, 0.8 g LDS (lithium dodecyl sulfate), 0.006 g EDTA (ethylenediaminetetraacetic acid), 4 g glycerol, 0.75 ml SERVA Blue G250 (1 % solution) and 0.25 ml phenol red (1% solution) and 10 X DTT (dithiothreitol) (Sigma-Aldrich) was added to each sample before heating at 70°C for 10 min (in cases where DTT was omitted, samples were heated for 5 min). Samples were then loaded onto a 4-12 % Bis-Tris NuPAGE gel (Invitrogen) and the gel run at 120 V for 1.5 hours. Protein was transferred to nitrocellulose membranes using the wet transfer Mini-Trans Blot Cell (Bio-Rad). The membrane was then washed with PBS and stained with Ponceau Red to allow visualisation of transferred protein. The membrane was then blocked for 1h with blocking solution (5 % western blotting grade blocker (BIO-RAD) and 1 % BSA (Sigma-Aldrich) in 1 X TBS-T) (described below). Following this the blots were probed with rabbit monoclonal anti-P-Trk (Cell Signaling, mAb #4621) diluted 1:4000, goat polyclonal anti-TrkB (R&D Systems) diluted 1:2000, goat polyclonal anti-TrkC (R&D Systems) diluted 1:2000, mouse monoclonal anti-synaptophysin (Sigma-Aldrich S5768) diluted 1:2000 or anti-arc mouse monoclonal (Santa Cruz sc – 17839). The primary antibody was diluted in blocking solution, and incubation was overnight at 4 °C.

The membranes were then washed 3 times with 20 ml TBS-T (10 X TBS (10 X TBS – 25 mM Tris Base, 1.37 mM NaCl, 2.6 mM KCl dissolved in ddH2O. TBS-T: 1 X TBS & 1% Tween-20 (Sigma-Aldrich) for 10 min per wash on a rocker and incubated with the secondary antibody (1:2000) in blocking solution for 1 hour. The primary antibodies were detected with donkey anti-Rabbit HRP-conjugated, anti-goat HRP-conjugated and anti-Mouse HRP-conjugated secondary antibodies (Promega). The membrane was developed using the WesternBright ECL Kit (Advansta). Blots were visualized with the Image Lab software and the Universal Hood III camera system (BIO-RAD). Densitometry analysis of the bands with ImageJ was used to calculate the intensity of the signal for each band. The quality of protein transfer was checked in every case by staining the membranes with Ponceau Red. Normalisation in all cases was deemed acceptable based Ponceau staining as the volume as well as the protein concentrations were found to be consistent between samples (see above). Synaptophysin was therefore not routinely used as a normalisation control.

### Small Angle X-ray scattering

Small angle X-ray scattering (SAXS) data were collected with in-line size exclusion chromatography (SEC-SAXS) at Diamond B21. Briefly, 10 mg/ml ZAB in Tris buffered saline was injected onto a Superdex 200 Increase column and data were collected across the elution volume. Data were analyzed, the buffer background scattering subtracted, and the peak data merged using the ScÅtterIV software(*80*). Ab-initio envelopes were created using DAMMIN (Dummy Atom Model Minimization) (*81*). Reciprocal space comparison of raw data and models was generated using CRYSOL (*82*). Data were plotted using SASPLOT (*82*).

## Supporting information

Supplemental Images

## Acknowledgments

The authors would like to acknowledge support from The Francis Crick Institute Structural Biology, Proteomics, and Chemical Biology STPs, particularly Dr Andy Purkiss, Dr Laura Masino, Dr Simone Kunzelman and Dr Dhira Joshi, and support from Diamond Light source Beamline Staff on beamlines I04, I04-1 and I24. AlphaFold3 was run using the Alphastream web interface implemented at the Francis Crick Institute by Dr Yew-Mun Yip.

## Funding

N.Q.M acknowledges that this work was supported by the Francis Crick Institute, which receives its core funding from Cancer Research UK (CC2068), the UK Medical Research Council (CC2068) and the Wellcome Trust (CC2068). This work was also funded by the Academy of Medical Sciences Springboard Award [SBF0010\1058] (C.J.P.) and the Biotechnology and Biological Sciences Research Council Doctoral Training Programme [BB/T0083690/1] (R.T.D.), and Barts Charity [G-002890] (P.C.A) Additional support was provided by Zebra Biologics Inc., and by NIH grant R41EY032011 (to P.S.D.).

## Author contributions

Conceptualization: DCB, YAB, RML, PSD, NQM. Methodology: DCB, CJP, SM, HN, PCA Investigation: DCB, SA, RTD, SM, HN, PCA Visualization: DCB, RTD, CJP,

Funding acquisition: NQM, YAB, RML, PSD, CJP, PCA Project administration: DCB, NQM

Supervision: NQM, CJP, YAB, RML, PSD Writing – original draft: DCB

Writing – review & editing: DCB, SA, RTD, SM, HN, YAB, PCA, CJP, PSD, RML, NQM

## Diversity, equity, ethics, and inclusion

The Francis Crick Institute believes that diversity drives scientific excellence and works to build an inclusive culture in which everyone can thrive. This paper has a diverse authorship in terms of gender and race and in keeping with our dedication to the UK Technician Commitment authorship, technicians and technical specialists have been given authorship status when this is in keeping with COPE guidelines and the Crick’s authorship policy, where authorship has not been granted contributions have been formally acknowledged.

## Competing interests

This work was in part supported by Zebra Biologics Inc. P.S.D. and R.M.L. are affiliated with Zebra Biologics Inc.

## Data, code, and materials availability

Map and models for the structures determined in this study are available at the Protein DataBank, under the accession numbers in table S1. All other data can be made available by request from the corresponding author.

