## Supplemental Images for "Discovery of a sulfotyrosine-motif in the human TrkB extracellular domain required for agonist activation"

**The PDF file includes:**

Figs. S1 to S6
Tables S1 to S2

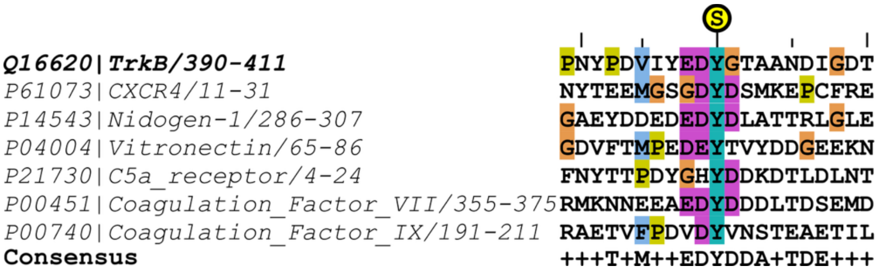


**FigS1**

**Similarity of the TrkB sTyr400 motif to known sTyr-containing motifs found in selected human proteins.** Uniprot identifiers and sequence extent are indicated.

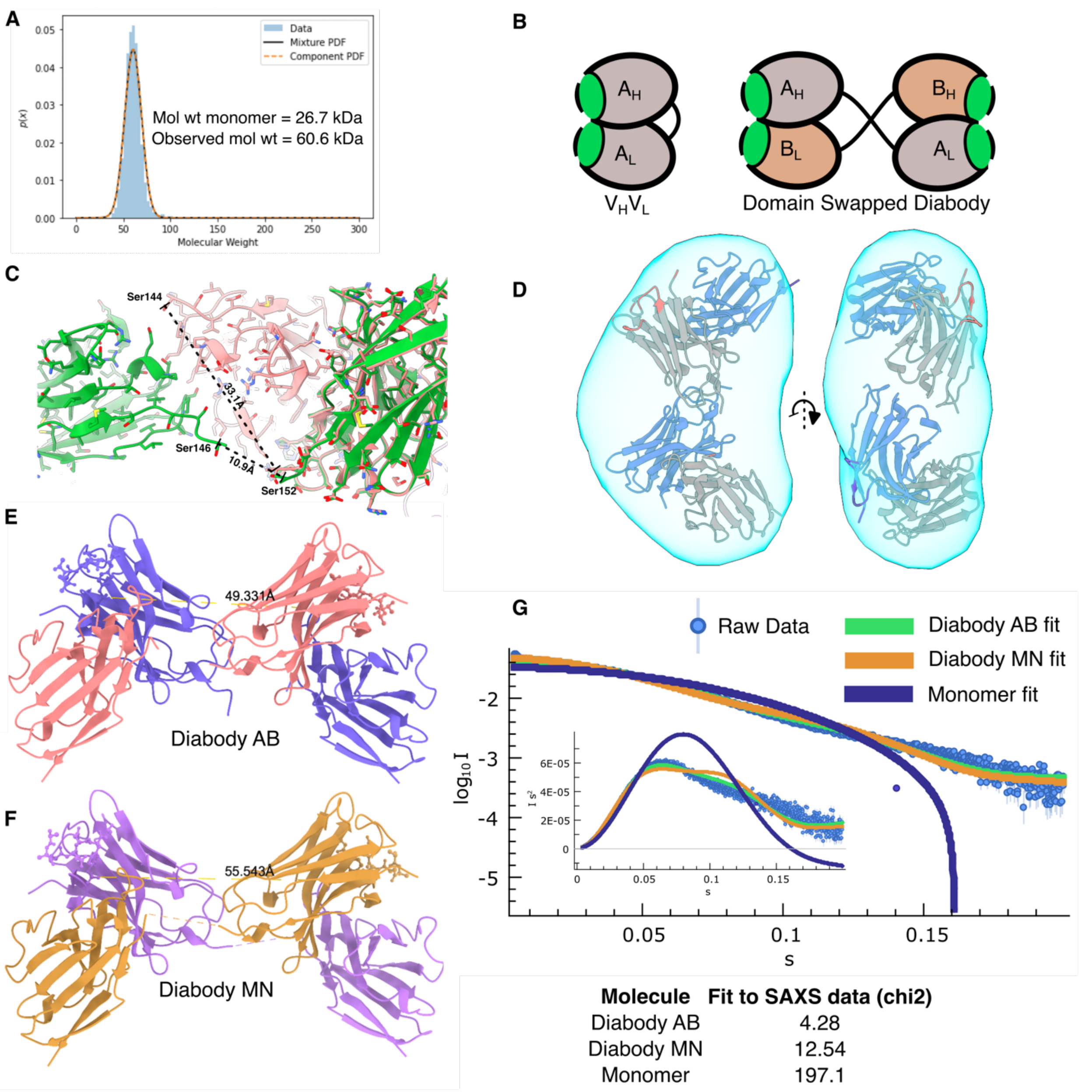


**Fig. S2.**

**Crystallographic and in solution evidence that ZAB85 adopts a dimeric state consistent with a diabody.** (**A**) Mass photometry analysis of ZAB85 is consistent with a dimeric oligomeric state even at low nanomolar concentrations, (**B**) Schematic of a monomeric compared to a domain-swapped diabody influenced by connecting linker length. (**C**) The disordered linker distance in the domain swapped, P1 model (green, missing residues 147-151, 10.9 Å) is plausible, whereas the monomeric configuration (pink, missing residues 145-151, 33.1 Å) is not. (**D**) Ab-initio SAXS DAMMIN envelope is consistent with a diabody configuration. (**E,F**) the two independent diabodies observed from the P1 cell processed crystal structure. The sTyr-sTyr binding site distances are indicated. (**G**) Diabody AB is most consistent with the SAXS data, although the Kratky plot (inset) is indicative of some flexibility.

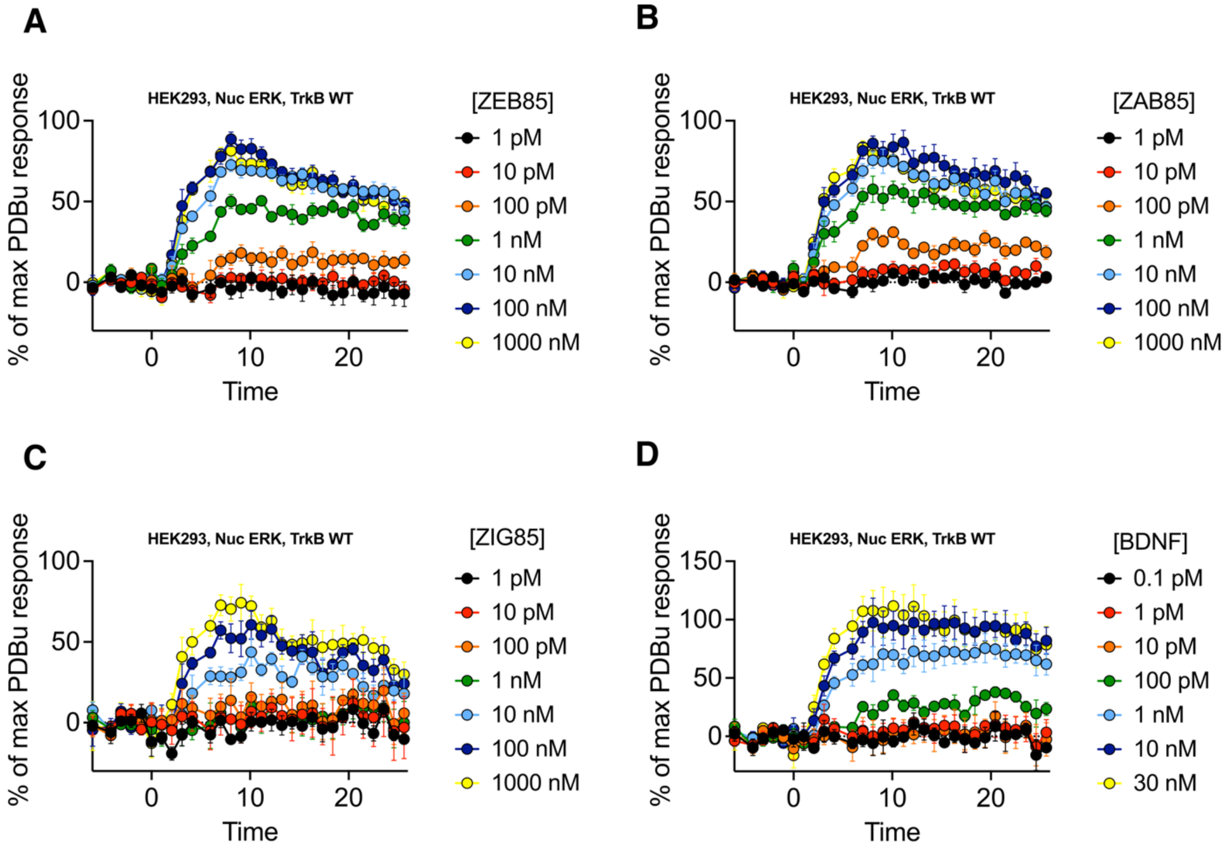


**Fig. S3:**

**Real-time ERK signalling in response to ZEB85, ZAB85, ZIG85 or BDNF' and** Human TrkB expressed in HEK293 cells alongside a nuclear ERK FRET sensor were stimulated with ZEB85 (A), ZAB85 (B), ZIG85 (C) or BDNF (D). Data from 4-5 independent replicates with triplicate wells. Mean ± SEM.

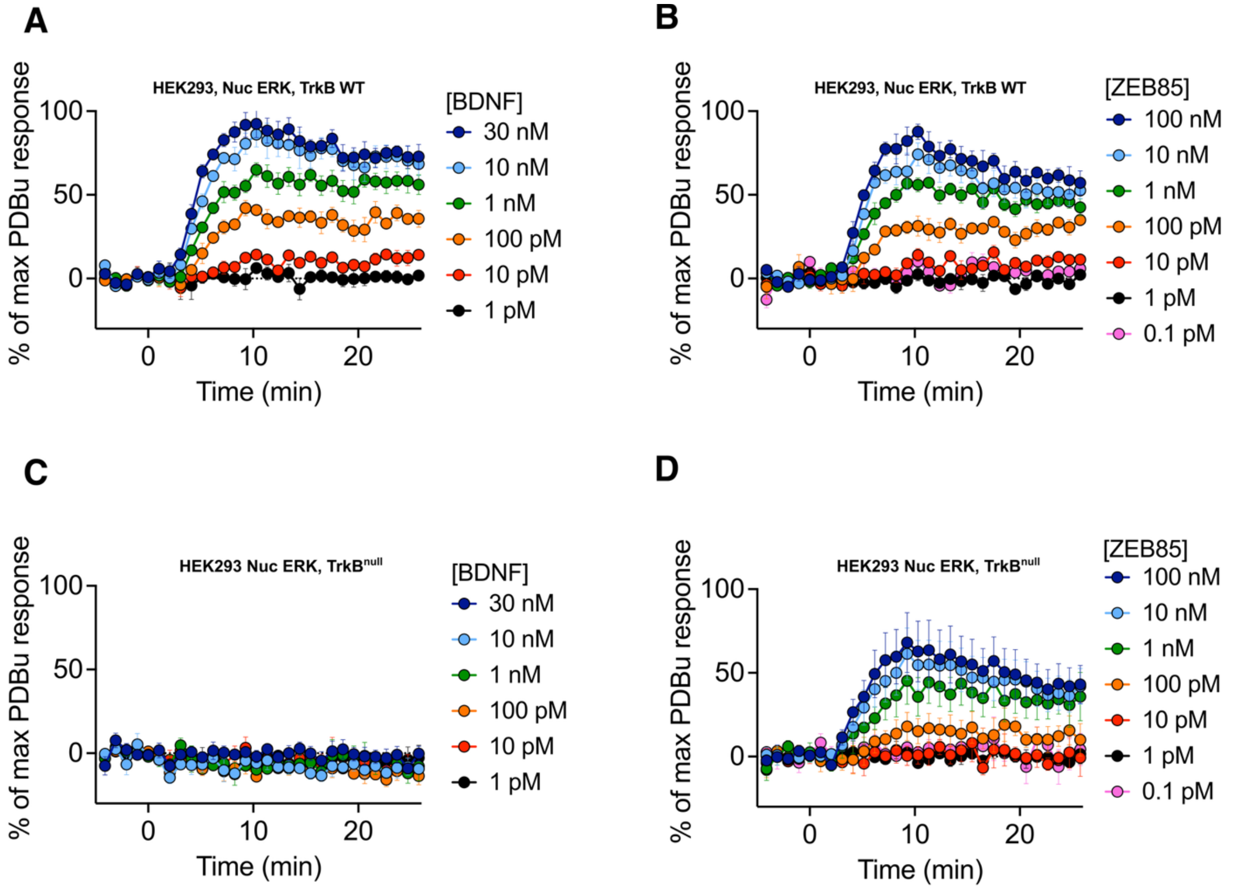


**Fig. S4**

**TrkB$^{null}$ disrupted BDNF-induced ERK signalling, but not ZEB85-induced responses.** Comparison of ERK responses to TrkB WT (A,B) or TrkB$^{null}$ (D298R/H299E/M379R) (C,D) in response to increasing concentrations of BDNF or ZEB85. Data from 4-5 independent replicates with triplicate wells. Mean ± SEM.

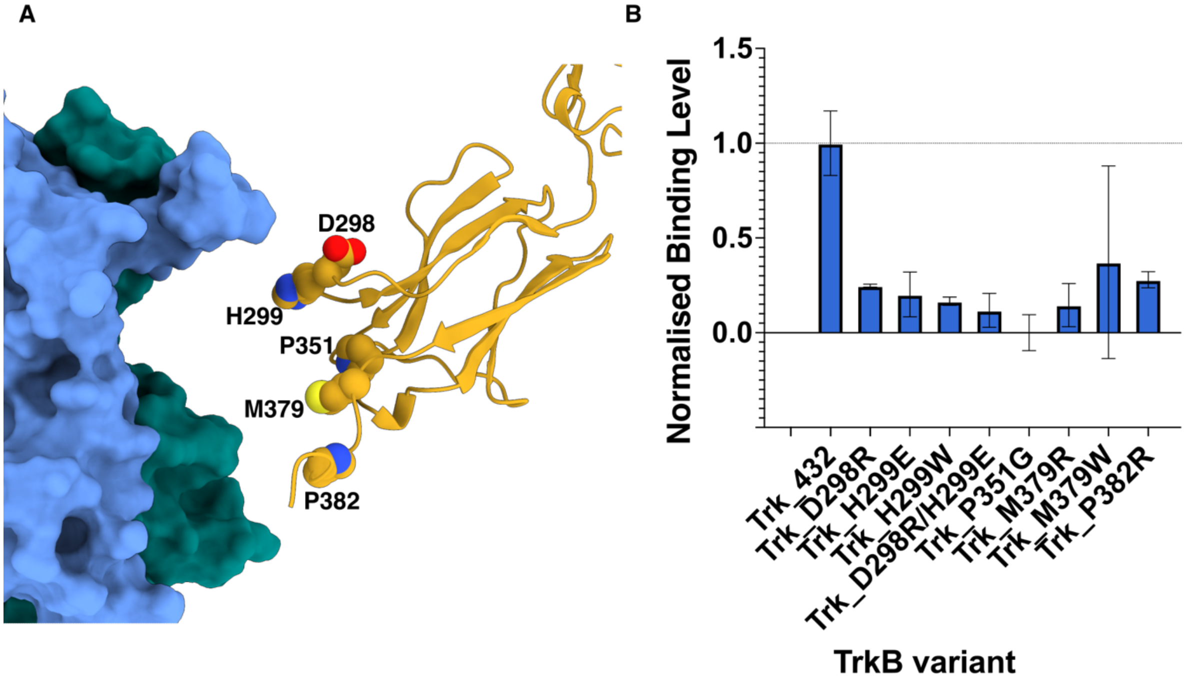


**Fig. S5.**

**Design of a BDNF-binding deficient TrkB mutant, TrkB$^{null}$.** (**A**) The sites of chosen mutations mapped onto our TrkB$^{383}$:BDNF$^{mat}$ structure, (interface opened for clarity) (**B**) SPR binding analysis of TrkB variants. Mean of 3 experiments ± S.D. Values are normalized to wild type TrkB$^{432}$.

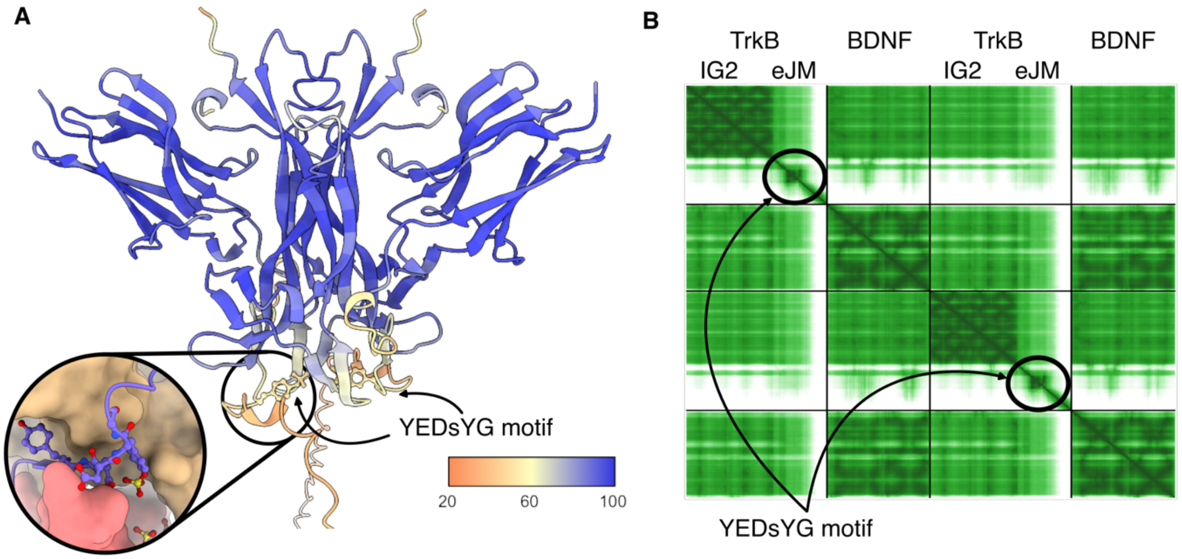


**Fig. S6.**

**Alphafold 3 models of the TrkB/BDNF interface suggest a role for the residues 383 to 405 of TrkB eJM in binding BDNF**. (**A**) The AlphaFold 3 model presented in Fig 3L and 3M coloured by pLDDT. Blue corresponds to high confidence, red corresponds to low confidence. Inset, shows a close-up of the predicted position of YEDsYG, occluded by BDNF. (**B**) Predicted aligned error plot for the same model. As before, the YEDsYG motif is highlighted.

**Table S1.**

| | ZEB85 VHL Apo<br>PDB: 30PO | ZEB85 VHL Bound<br>PDB: 30PP | TrkB:BDNF complex<br>PDB: 30PU |
|---|---|---|---|
| Wavelength | 0.9795 Å | 0.6199 Å | 0.9796 Å |
| Resolution range | 55.28 - 1.88<br>(1.94 - 1.88) | 42.59 - 2.06<br>(2.13 - 2.056) | 64.19 - 3.0<br>(3.11 - 3.0) |
| Space group | P 64 2 2 | P 64 2 2 | C 1 2 1 |
| Unit cell | a=b=110.56<br>c= 184.30<br>90 90 120 | a=b= 110.96<br>c= 184.01<br>90 90 120 | a= 56.878<br>b=179.73 c=183.45<br>90 90.04 90 |
| Total reflections | 6541514 (609132) | 3194219 (323105) | 506973 (45341) |
| Unique reflections | 54723 (5352) | 42061 (4109) | 36805 (3618) |
| Multiplicity | 119.5 (113.8) | 75.9 (78.6) | 13.8 (12.5) |
| Completeness (%) | 99.93 (100.00) | 99.72 (99.47) | 69.96 (25.06)<br>(*STARANISO processed*) |
| Mean I/sigma(I) | 21.55 (0.31) | 11.80 (0.59) | 6.35 (0.09) |
| Wilson B-factor | 48.46 | 47.87 | 107.63 |
| R-pim | 0.01209 (0.6188) | 0.02689 (0.3572) | 0.05366 (4.401) |
| CC1/2 | 1 (0.425) | 1 (0.316) | 0.982 (0.179) |
| CC* | 1 (0.772) | 1 (0.693) | 0.995 (0.552) |
| Reflections used in refinement | 54687 (5352) | 42039 (4109) | 25738 (912) |
| Reflections used for R-free | 2669 (270) | 2102 (201) | 1349 (42) |
| R-work | 0.1934 (0.3914) | 0.2136 (0.3887) | 0.2531 (0.4131) |
| R-free | 0.2317 (0.4213) | 0.2494 (0.3766) | 0.2835 (0.4426) |
| Number of non-hydrogen atoms | 3803 | 3844 | 7409 |
| macromolecules | 3550 | 3714 | 7311 |
| ligands | 94 | 71 | 98 |
| solvent | 207 | 101 | 0 |
| Protein residues | 459 | 478 | 936 |
| RMS(bonds) | 0.012 | 0.003 | 0.003 |
| RMS(angles) | 1.09 | 0.61 | 0.61 |
| Ramachandran favored (%) | 96.45 | 96.09 | 92.53 |
| Ramachandran allowed (%) | 3.55 | 3.91 | 7.14 |
| Ramachandran outliers (%) | 0.00 | 0.00 | 0.32 |
| Rotamer outliers (%) | 0.26 | 0.25 | 0.84 |
| Clashscore | 4.11 | 2.60 | 3.83 |
| Average B-factor | 61.09 | 63.62 | 106.01 |
| macromolecules | 61.02 | 63.82 | 105.38 |
| ligands | 69.46 | 68.22 | 152.85 |
| solvent | 60.38 | 55.20 | N/A |

**X-ray Crystallographic data and refinement statistics**

Statistics for the highest-resolution shell are shown in parentheses.

**Table S2**

| Immobilised Partner | Immobilisation method | Analyte | $K_D$* |
|---|---|---|---|
| **TrkB_383** | Amine Coupling | ZEB85 | N.B |
| **TrkB_432** | " | ZEB85 | 0.24 nM ± 0.08 nM |
| **TrkB_383-432 (Y400)** | Biotin/Streptavidin | ZEB85 | 2.34 μM ± 0.91 μM |
| **TrkB_393-405 (Y400)** | " | ZEB85 | N.B |
| **TrkB_393-405 (sY400)** | " | ZEB85 | 0.52 nM ± 0.71 nM |
| **BDNF** | Amine Coupling | TrkB_432 | 9.8 nM ± 6.24 nM |
| **"** | " | TrkB_421 | 12.1 nM ± 1.15 nM |
| **"** | " | TrkB_412 | 3.8 nM ± 0.49 nM |
| **"** | " | TrkB_402 | 9.1 nM ± 0.19 nM |
| **"** | " | TrkB_393 | 37.7 nM ± 32.5 nM |
| **"** | " | TrkB_383 | 35.7 nM ± 18.4 nM |
| **"** | " | TrkB_432_D298R_H299E | N.B |
| **"** | " | TrkB_432_Y400F | 15.3 nM ± 6.6 nM |

**Summary of Surface Plasmon resonance data**

*Geometric mean ± 0.5x range, all SPR experiment n=3 or greater.

N.B = no binding detected.
